# Harmonizing Inter-Site Differences in T1-Weighted Images Using CycleGAN

**DOI:** 10.1101/2025.04.07.646944

**Authors:** Masaaki Shimizu, Yuichi Yamashita, Hiroyuki Yamaguchi, Naoya Oishi, Hidehiko Takahashi, Genichi Sugihara

## Abstract

**Introduction:** When neuroimaging studies using magnetic resonance imaging (MRI) are conducted across multiple centers, they often encounter inter-site differences in MRI equipment and protocols leading to biases and confounding effects in MRI measurements. There are existing techniques for correcting these site effects, i.e., harmonization, but they have limitations, including the need for preprocessing of MRI data, which involves processes such as spatial normalization. Deep learning-based methods have emerged as potential alternatives that can handle site effects without the need for preprocessing steps. In this study, we propose a novel method based on the generative adversarial network (GAN) framework, CycleGAN, that effectively addresses inter-site differences in T1-weighted images with minimal preprocessing requirements. We compare the harmonization efficacy of CycleGAN with that of the commonly used method ComBat.

**Methods:** We trained the proposed CycleGAN method and the comparative ComBat method using data from 40 subjects at each of two sites. To evaluate the effectiveness of the two methods, we used data from nine subjects who underwent imaging at both sites. We assessed harmonization performance at the image level using the structural similarity index (SSIM) and peak signal-to-noise ratio (PSNR). Additionally, we evaluated harmonization results at the feature level by analyzing regional cortical thickness and volume data. Cohen’s *d* was employed to quantify the differences between feature values.

**Results:** At the image level, the ComBat method decreased the median baseline SSIM value from 0.86 (interquartile range [IQR], 0.02) to 0.84 (IQR, 0.02), whereas the proposed CycleGAN method maintained the SSIM value at 0.86 (IQR, 0.02). For PSNR, the baseline value was 18.33 (IQR, 1.78), which decreased to 15.30 (IQR, 2.20) after applying ComBat, but increased to 19.58 (IQR, 3.12) with the proposed CycleGAN method. These findings indicate that CycleGAN preserved the structural and signal similarity of the images. At the feature level, the effect size for cortical thickness decreased from 0.97 (IQR, 1.79) to 0.91 (IQR, 1.54) after applying ComBat, whereas the proposed CycleGAN method yielded an effect size of 1.05 (IQR, 1.14). For cortical volume, the effect size decreased from 0.95 (IQR, 1.78) to 0.69 (IQR, 1.00) after applying ComBat, and decreased to 0.88 (IQR, 0.74) with the CycleGAN method. Compared with baseline, Cohen’s *d* was significantly lower with both ComBat (p = 0.000002) and CycleGAN (p = 0.028) with no significant difference between the two methods, indicating similar performance of the two methods under the study conditions.

**Conclusion:** The results underscore the ability of CycleGAN to harmonize data without explicit normalization and emphasize the potential impact of the normalization process on harmonization procedures.

Our findings suggest that CycleGAN holds promise as a harmonization technique in multi-site neuroimaging studies.

## Introduction

Neuroimaging techniques, particularly magnetic resonance imaging (MRI), have advanced our understanding of the neural basis of neurological and psychiatric disorders and have provided biomarkers (Cortese et al., 2021; Kraguljac et al., 2021; Zackova et al., 2021; Zhang et al., 2018). However, the heterogeneity and complexity of these disorders pose significant challenges for detecting clinically meaningful alterations in MRI measurements. One of the major sources of heterogeneity is high intersubject variability in both pathology and symptoms, which can lead to large variability in MRI measurements across patients with the same disorder (Toal et al., 2010; Arnone et al., 2009; Miranda et al., 2021; Zhang et al., 2015; Zhuo et al., 2019). Additionally, subtle alterations in MRI measurements can be difficult to detect, even with large sample sizes.

The median sample size in neuroimaging research is reported to be 25, but it has been suggested that a sample size of over a thousand is preferable when factors such as reproducibility are considered (Marek et al., 2022; Yamashita et al., 2019). To address these challenges, international frameworks for sharing data across multiple sites have been initiated, such as the Strategic Research Program for Brain Science (SRPBS) (Tanaka et al., 2021) and the Autism Brain Imaging Data Exchange (ABIDE) (Martino et al., 2014; Martino et al., 2017). However, multi-site data also contain inter-site differences due to differences in MRI equipment and imaging protocols, which can lead to systematic biases and confounds in the MRI measurements (Takao et al., 2011; Han et al., 2006; Takao et al., 2013; Yamashita et al., 2019). Therefore, the removal of inter-site differences in neuroimaging data, i.e., harmonization, is an important challenge when elucidating the pathological mechanisms of disorders and identifying reliable biomarkers.

Thus far, various techniques have been used for harmonization. The ComBat method (Fortin et al., 2017; Johnson, Li, and Rabinovic, 2007) and the traveling subject (TS) method (Yamashita et al., 2019) are currently popular because of their efficacy in correcting site effects. Both techniques employ an extension of the general linear model (GLM) that includes site variables as categorical variables, with these being used to estimate and eliminate site effects. Moreover, ComBat models scaling factors specific to each site and improves parameter estimation for smaller samples by utilizing an empirical Bayes method. An additional benefit of this method is that it can be used for retrospective study. TS method is a bias estimation technique, employing traveling subjects who undergo scanning procedures at distinct sites. Because the subjects are consistent across all sites, TS method eliminates the need to select and include covariates, such as age and gender, in the model.

Despite the effectiveness of the GLM-based methods for correcting site effects, several issues have yet to be addressed. First, the current methods need to be applied after extraction of features such as cortical thickness and functional connectivity, because these feature values can vary according to software, the computer operating system, and system memory (Gronenschild et al. 2012; Hedges et al. 2021; Chepkoech et al. 2016). Furthermore, precise estimation of site effects becomes difficult when site and other covariates such as age and sex are correlated. Second, because existing GLM-based methods are based on linear models, they cannot eliminate nonlinear effects. Nonlinear trends, such as changes in cortical volume, require generalized additive models that account for nonlinearities in the covariates (Pomponio et al., 2020). These issues suggest that site effects may be more complex than previously thought, thus necessitating the development of novel harmonization methods.

Recently, deep learning-based methods have emerged as potential alternatives to overcome the limitations of GLM-based approaches. These methods use MRI data as input, do not require feature extraction, and can handle nonlinear site effects; furthermore, they have been shown to effectively remove site effects in MRI data (Zhao et al., 2019; Dinsdale et al., 2020; Moyer et al., 2020; Yamaguchi et al., 2021). However, previous deep learning-based approaches, such as variational autoencoder and U-Net (Moyer et al., 2020; Dewey et al., 2019), required several preprocessing steps prior to harmonization, including skull-stripping and spatial normalization.

Because these steps may be sensitive to site effects, it is desirable to minimize the preprocessing needed to apply the harmonization method.

In this study, we propose a novel harmonization method using a deep learning-based approach, specifically, the generative adversarial network (GAN). GAN is a machine learning algorithm comprised of two neural nets, a generator and a discriminator, which interact in an adversarial manner. During the training process, the generator generates fake images that the discriminator then evaluates. The discriminator is trained to distinguish the real images from the fake ones, and its feedback is used to improve the generator’s ability to produce more realistic data (Figure 1). We employ cycle GAN (Zhu et al., 2017), an extension of GAN, for image-to-image translation tasks, where it eliminated the need for paired training data. Cycle GAN utilizes two generator-discriminator pairs, one to transform images from domain A to domain B and another to transform images from domain B to domain A. Cycle GAN has been effective in various image-to-image translation tasks, such as transforming horses into zebras and converting summer scenes into winter ones. Typically, these deep learning approaches require a substantial amount of training data to prevent overfitting while training the model. However, the acquisition of a large dataset of MRI images in a short period is not feasible for every facility. Thus, our research focuses on a limited dataset of images from 40 individuals, demonstrating that our method can achieve a significant level of accuracy even with this scale of training data and minimal preprocessing. Additionally, whereas deep learning methods generally rely on large datasets to prevent overfitting, our approach shows that overfitting can be mitigated to a considerable extent even with a small number of data points. This is evidenced by using a unique dataset of “Traveling Subjects” individuals (Yamashita et al., 2019) who have been imaged at multiple facilities. This highlights the importance and potential of applying deep learning techniques in scenarios where data availability is limited and this aspect, not explored in similar previous research, represents a significant contribution of our study and is a point we wish to strongly emphasize.

**Fig. 1.**
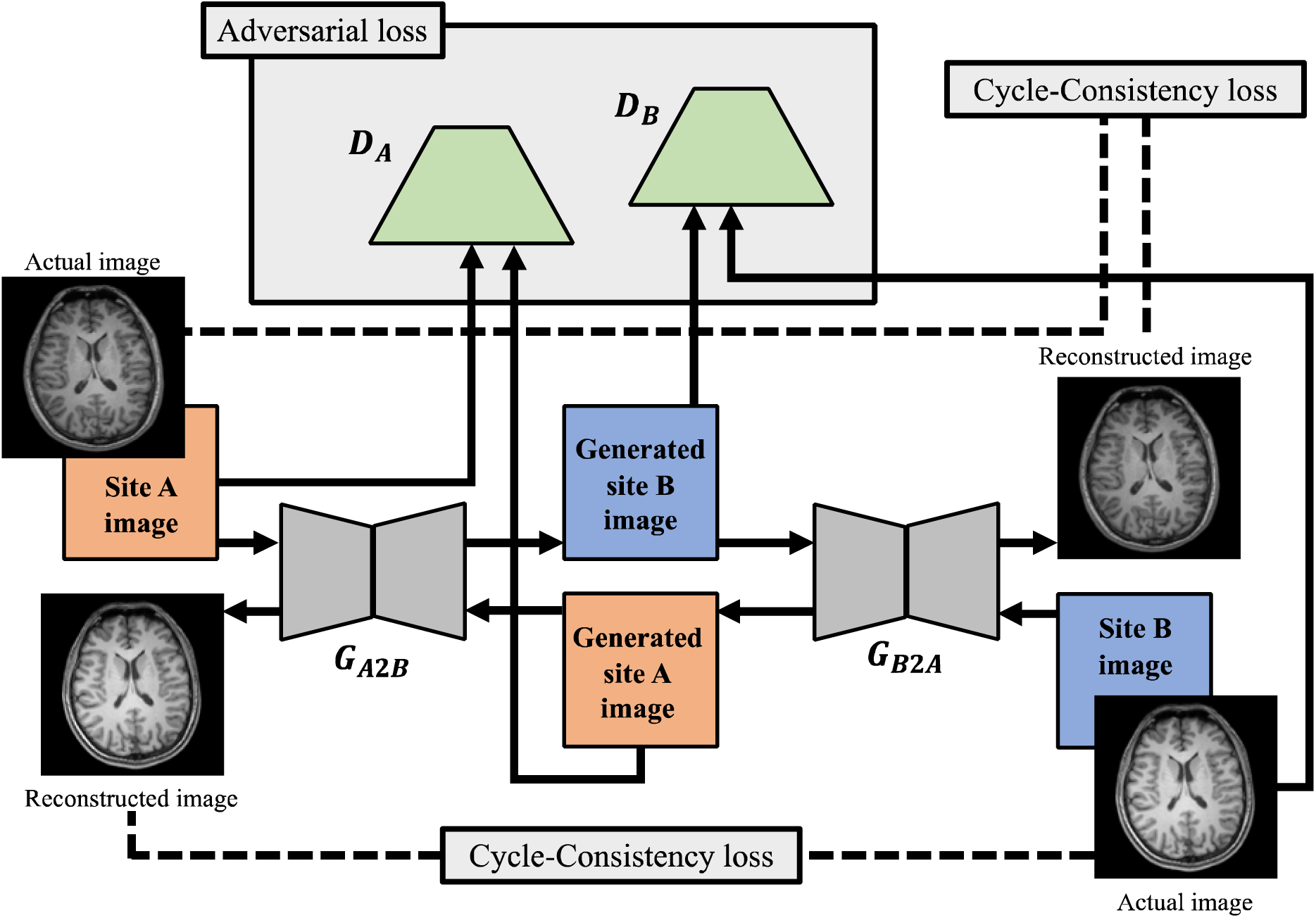
The fundamental architecture of CycleGAN.

## Methods

Ethical approval for this study was obtained from the Ethics Committee at Tokyo Medical and Dental University (Approval Number M2019-080).

### 2.1 Dataset

T1-weighted images from the SRPBS dataset (Tanaka et al., 2021) from a brain image database in Japan and acquired at Kyoto University (KUT) and Showa University (SWA) were used as training data, with 40 healthy subjects per institution. We used the TS dataset as test data (Yamashita et al., 2019). Both Dataset were obtained from the DecNef Project Brain Data Repository (https://bicr-resource.atr.jp/srpbsts/) gathered by a consortium as part of the SRPBS supported by the Japanese Advanced Research and Development Programs for Medical Innovation (AMED). This dataset is a collection of images from nine healthy subjects that were acquired at each site. The TS dataset contains data acquired at KUT and SWA with the same imaging parameters as used for the SRPBS data. Demographic data of the subjects in the training and test sets are summarized in Table 1.

**Table 1.** Characteristics of the sample.

| Training data |  |  |
| --- | --- | --- |
|  | SWA | KUT |
| n (subjects) | 40 | 40 |
| Age (year) mean (SD) | 30.9 (8.8) | 33.5 (9.5) |
| Gender (male) | 25 | 21 |
| Handedness (right) | 38 | 39 |
| Traveling subject data |  |  |
|  | SWA | KUT |
| n (subjects) | 9 | 9 |
| Age (year) mean (SD) | 27.0 (2.6) | 27.0 (2.6) |
| Gender (male) | 9 | 9 |

### 2.2 MRI image acquisition and preprocessing

At SWA, T1-weighted images were acquired with a 3T MRI scanner (Siemens Tim Trio) using a 32-channel head coil. The imaging parameters included: repetition time (TR) = 2000 ms, echo time (TE) = 3.4 ms, image dimensions = 240 × 256 × 208, flip angle = 8°, voxel size = 1 × 1 × 1 mm, and phase encoding direction = posterior–anterior. At KUT, T1-weighted images were acquired with a 3T MRI scanner (Siemens Verio) using a 12-channel head coil. The imaging parameters included: TR = 2300 ms, TE = 2.98 ms, image dimensions = 240 × 256 × 256, flip angle = 9°, voxel size = 0.9375 × 0.9375 × 1, and phase encoding direction = posterior– anterior. Because the voxel size and image dimensions differed between the sites, we conducted minimal preprocessing involving transforming the voxel size to 1 × 1 × 1 mm and the dimensions to 256 × 256 × 256 using <u>FreeSurfer software</u> functions. No other preprocessing was performed.

### 2.3 Cycle GAN

We used the CycleGAN architecture, which is a model that extends GAN, for the proposed harmonization method (Goodfellow et al., 2014). CycleGAN contains two generators and two discriminators. The two generators were each trained to transform images acquired from site A into those representative of site B (G_A2B_), and vice versa (G_B2A_). The two discriminators also learned to correctly discriminate whether the input image was an image generated by a generator or an actual image.

Three losses are determined for the objective of optimizing the model, adversarial loss, cycle-consistency loss, and identity loss.

(a) Adversarial loss: This loss will be small if the generator makes the discriminator recognize the generated images as the actual image. Simultaneously, the loss will also be small if the discriminator correctly distinguishes between the generated and actual images (hereafter, actual site A images are denoted as *a* and actual site B images are denoted as *b*). *G_A2B_* and *G_B2A_* represents the generator that converts input images into images acquired at site B and site A respectively. *D_A_* represents the discriminator that discriminates whether input images are actual site A images or are created by the generator. *D_B_* similarly represents the discriminator that discriminates whether input images are actual site B images or are created by the generator.

In this way, by setting up the described losses, the generator is trained to produce images that the discriminator judges as actual in the first term, while in the second term, the discriminator is trained to correctly classify actual images. Therefore, the relationship between these generators and discriminators is referred to as adversarial loss.

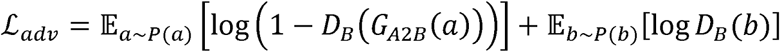

(b) Cycle-Consistency loss: The images generated from actual site A images by *G_A2B_(a)* are input to the generator *G_B2A_*. Then, the loss is set so that the difference between the resulting output *G_B2A_(G_A2B_(a))* and the original a is small.

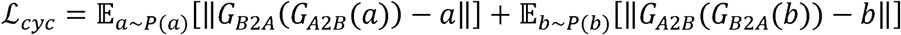

(c) Identity loss: This loss is defined to verify that the generated image is close to the actual site A image, even if the actual site A image is input generated by *G_B2A_*.

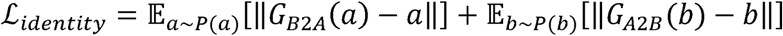

The data used for CycleGAN are described in Table 1. The TS data were used as the test dataset. Each 3D MRI images from the datasets were first split into axial 2D slices to prepare the data for processing with CycleGAN. Consequently, the input images were 256 × 256 two-dimensional images that were slices of three-dimensional brain scans acquired in the axial direction. The generators consist of three convolutional layers followed by nine stacked ResNet blocks then two more convolutional layers. The discriminators are made up of five convolutional layers. Both have input dimensions of 256 × 256. In addition, scaling was applied to adjust the dynamic range of the images after CycleGAN to 255. The details of architectures, their training and other information are presented in the Supplementary Materials.

### 2.4 ComBat

We used ComBat as a comparison method to evaluate the performance of our proposed CycleGAN method. The training data consisted of 40 images from the SRPBS dataset, acquired at KUT and SWA. Since ComBat necessitates that input images are spatially normalized, we performed the following procedures. First, the SRPBS and TS data were spatially normalized to the ICBM MNI152NLin2009cAsym template using the Computational Anatomy Toolbox (CAT12) (Gaser et al., 2022), saving the transformation parameters used. Next, the parameters of ComBat were set from the normalized training data. We then applied these parameters to the normalized TS data to eliminate estimated site effects from the images. Finally, the corrected images were transformed back to the individual brain space by multiplying them by the inverse transformation parameters. Subsequently, we measured the regional cortical thickness and volume of these images using FreeSurfer (version 7.1.1). A more detailed ComBat model can be found in the Supplementary material.

### 2.5 Evaluation and statistical analyses

In this study, we performed evaluations using both image measurements and measurements of extracted brain features. For the evaluation of the T1-weighted images, we used the structural similarity index (SSIM) and peak-signal to noise ratio (PSNR), which were previously used in similar studies (Zuo et al., 2021; Liu et al., 2021; Duffy et al., 2021). The SSIM is a metric representing the similarity between two image pairs, and is calculated using the following equation.

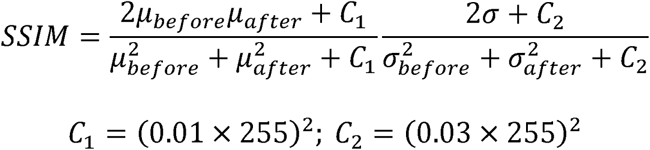

where “before” and “after” mean the images before harmonization and after harmonization. SSIM evaluates images by dividing them into small regions and then assessing each region. The variable “μ” represents the average pixel value in the small region, “σ” represents the standard deviation in the same region, and “C” is a constant used to avoid division by zero. SSIM is used to evaluate the structural similarity between subjects (images) and was calculated for each axial slice. Therefore, a higher SSIM implies that the structural similarity is better preserved. In this study, we expected that the structural similarity would be maintained before and after harmonization, so a higher value of this index was preferable.

PSNR is a metric that represents the ratio of the maximum pixel value of an image to the noise, and was calculated using the equation:

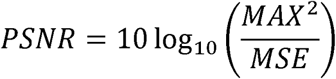

Here, we define noise as the mean squared error between the two images. “MAX” represents the maximum pixel value of the image, and “MSE” represents the mean squared error for each pixel. In MRI, an improvement in PSNR signifies an enhancement in the quality of the image. Higher PSNR indicates that more information is retained between the original and corrected images, making a higher value of this index preferable.

For feature-level evaluation, we used regional cortical thickness and volume. The average cortical thickness and volume were calculated using FreeSurfer (ver 7.1.1) for each of the 68 regions defined in the Desikan-Killiany atlas (Desikan et al., 2016). We used paired Cohen’s *d* as a metric for the feature value difference between the two conditions of each method. Cohen’s *d* is an effect size index representing the standardized difference in mean values. A larger Cohen’s *d* indicates larger difference in the mean values between the two groups; more concisely, a *d* value of 0 indicates that the two groups have identical means, while a *d* value of 1 indicates that the difference between the means is equal to one standard deviation. Cohen’s *d* was expressed using the following equation, where *i* is the number of subjects in each group and *x_i_* and *y_i_* are the variables for the *i*-th subject in each group. In this study, *x* and *y* correspond to features of the ROI in images from KUT and SWA.

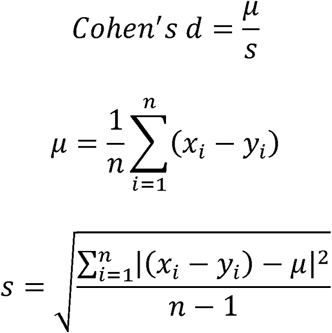

To quantify the site effect, we calculated the SSIM and PSNR between SWA and KUT images using the uncorrected test data (between actual SWA images and actual KUT images). These metrics were used as a baseline to assess the effectiveness of the proposed method. To evaluate the proposed method, we generated SWA images of the test data and compared the metrics between the generated images and the actual SWA data. Thus, we compared each evaluation index under three conditions: baseline, after applying ComBat, and after applying the proposed CycleGAN method. Pairwise Wilcoxon signed rank tests were used to compare the results. The statistical analyses were performed with <u>R</u>.

## Results

Figure 2 shows examples of original images and harmonized images. Qualitative comparisons reveal that the images harmonized by Cycle GAN have successfully corrected inter-site differences in comparison with the original images. Note that the CycleGAN-corrected images resemble the reference images because CycleGAN harmonizes images from one site to another by transforming them, whereas the ComBat-corrected images show both sites becoming more similar to each other because ComBat harmonizes inter-site differences by removing site-specific components from the images. In other words, in contrast to CycleGAN, which generates images from one site to another, ComBat produces harmonized images by removing site-specific components from images of each site.

**Fig. 2.**
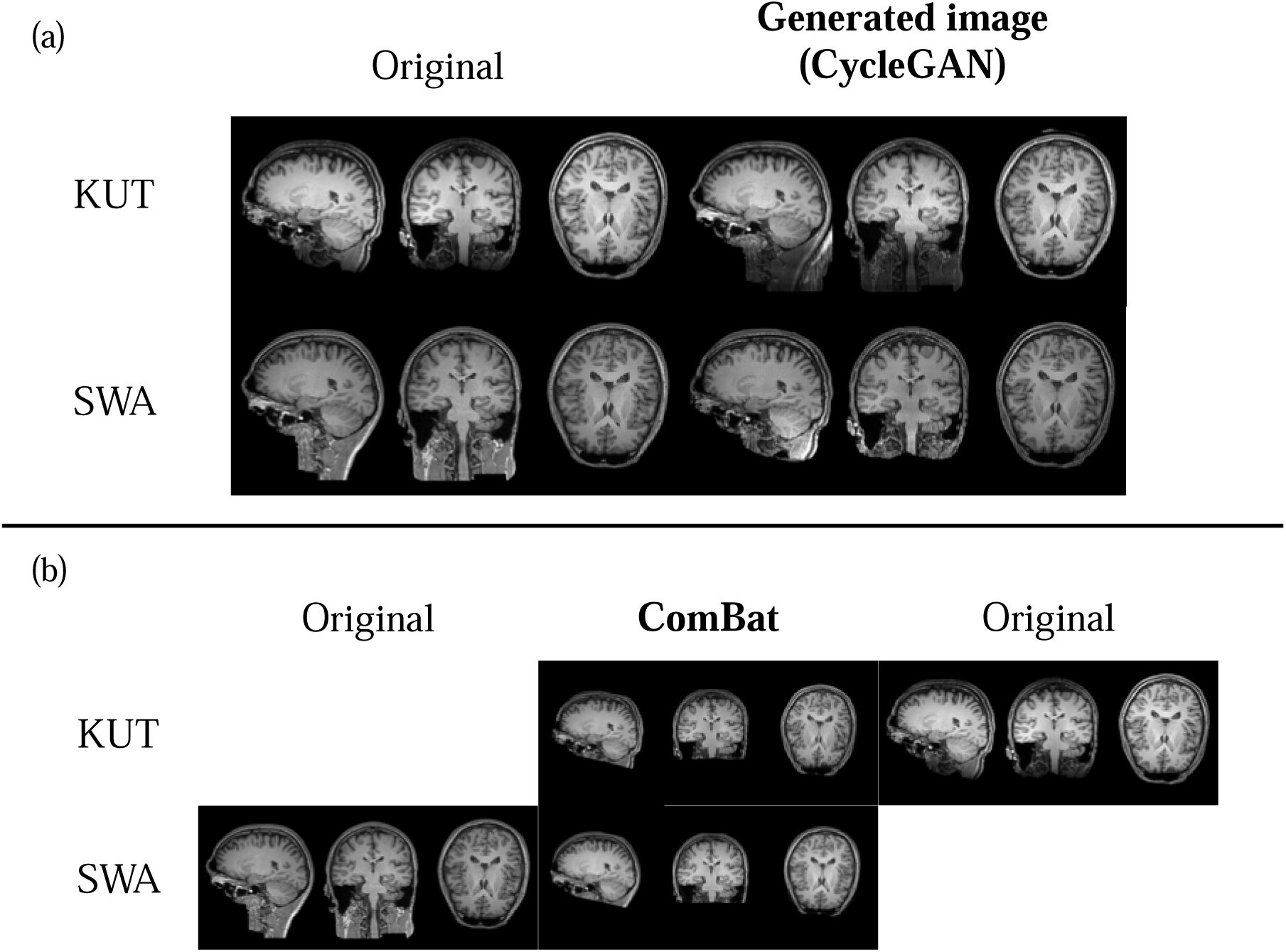
Examples of actual images. (a) (Top Left) TS data acquired at KUT. (Bottom Left) TS data acquired at SWA. (Top Right) Image generated by CycleGAN, KUT-type image generated by CycleGAN from input of SWA TS data. (Bottom Left) SWA-type image generated by CycleGAN from input of KUT TS data. (b) (Bottom Left) TS data acquired at SWA. (Top Center) Image obtained by applying ComBat to KUT TS data. (Bottom Center) Image obtained by applying ComBat to SWA TS data. (Bottom Right) TS data acquired at KUT.

The changes in SSIM and PSNR before and after applying ComBat and our proposed method are shown in Figure 3. The alterations in SSIM are presented in Figure 3a, and demonstrate the impact of both the ComBat and CycleGAN methods. The baseline median SSIM value was 0.86 (interquartile range [IQR] 0.02). After applying the ComBat method, this SSIM value decreased to 0.84 (IQR, 0.02), whereas application of our proposed CycleGAN method resulted in an SSIM SSIM values between the conditions (χ^2^(2) = 8.67, p = 0.013). The pairwise Wilcoxon signed value of 0.86 (IQR, 0.02). The Friedman test revealed statistically significant differences in rank test showed significant differences in SSIM between baseline and ComBat images (p = 0.008), as well as between ComBat and CycleGAN images (p = 0.008). Nevertheless, no significant difference in SSIM was observed between the baseline and CycleGAN conditions. These findings suggest a significant decrease in SSIM values after applying the ComBat method in comparison with both the baseline and CycleGAN.

**Fig. 3.**
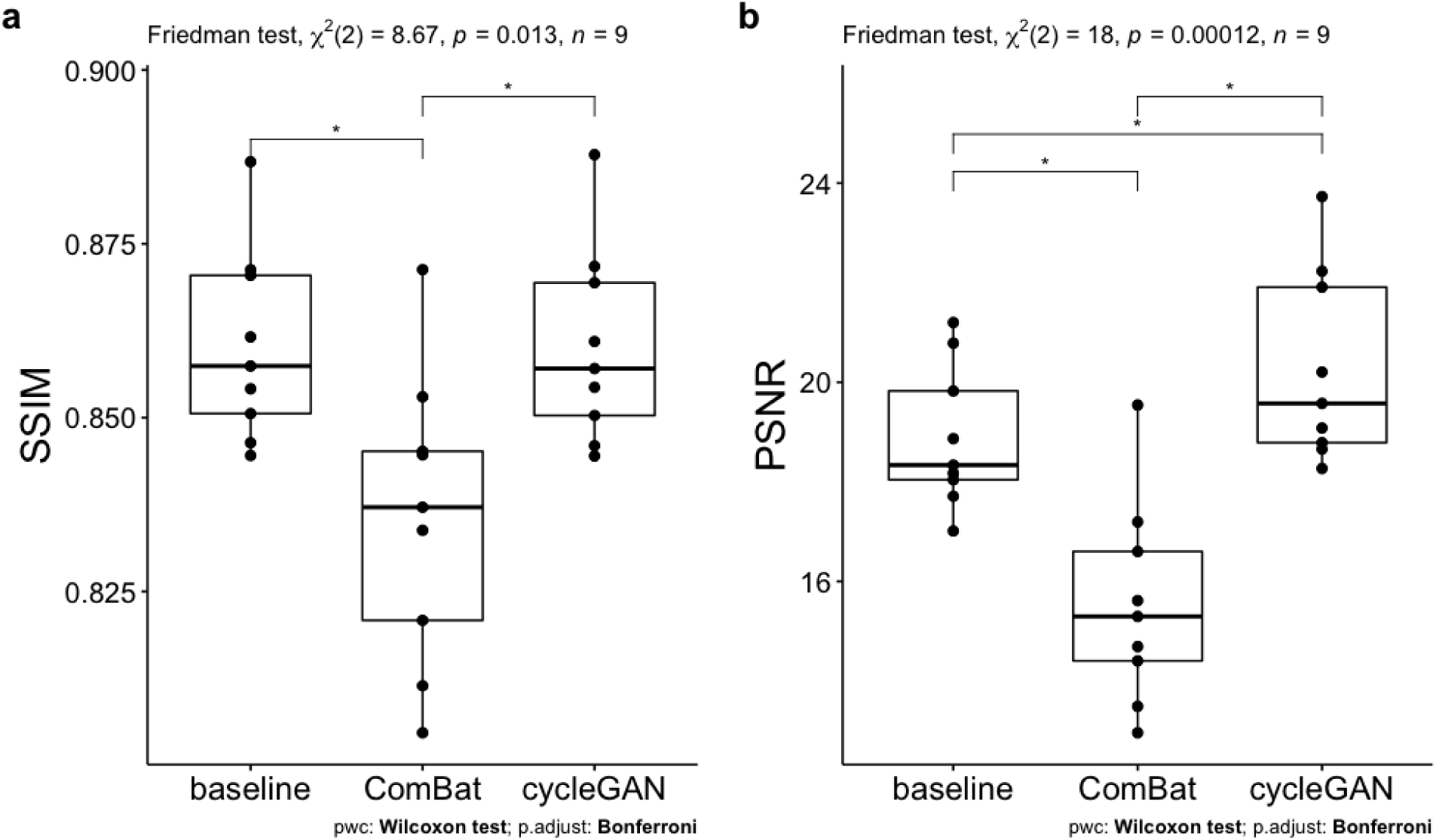
Results of SSIM (left) and PSNR (right) analyses for the three conditions of baseline, ComBat, and CycleGAN. Each dot corresponds to nine subjects. * = p < 0.05

The changes in PSNR before and after applying ComBat and CycleGAN are presented in Figure 3b. The median PSNR values were 18.33 (IQR, 1.78) for baseline, 15.30 (IQR, 2.20) after applying the ComBat method, and 19.58 (IQR, 3.12) after applying CycleGAN. The Friedman conditions (χ^2^(2) =18, p = 0.0012). Pairwise Wilcoxon signed rank tests between conditions test indicated a statistically significant difference in PSNR values between the different revealed statistically significant differences in PSNR between baseline and ComBat (p = 0.004), ComBat and CycleGAN (p = 0.004), and baseline and CycleGAN (p = 0.004). These results show that in comparison with baseline images, PSNR was significantly improved after applying our proposed method.

As in the procedure for SSIM and PSNR, Cohen’s *d* between the SWA and KUT images of the test data was used as the baseline for assessing the effects on extracted measurements. The results for cortical thickness are shown on the left side of Figure 4a. The median effect size at baseline was 0.97 (IQR, 1.79). After applying ComBat this was 0.91 (IQR, 1.54), and after applying CycleGAN it was 1.05 (IQR, 1.14). The Friedman test showed that Cohen’s *d* was not significantly different between the conditions ( (2) =1.68, p = 0.43). There was no significant difference in mean Cohen’s *d* between the baseline and either processing condition. However, after applying ComBat and CycleGAN there was less variability and lower mean values.

**Fig. 4.**
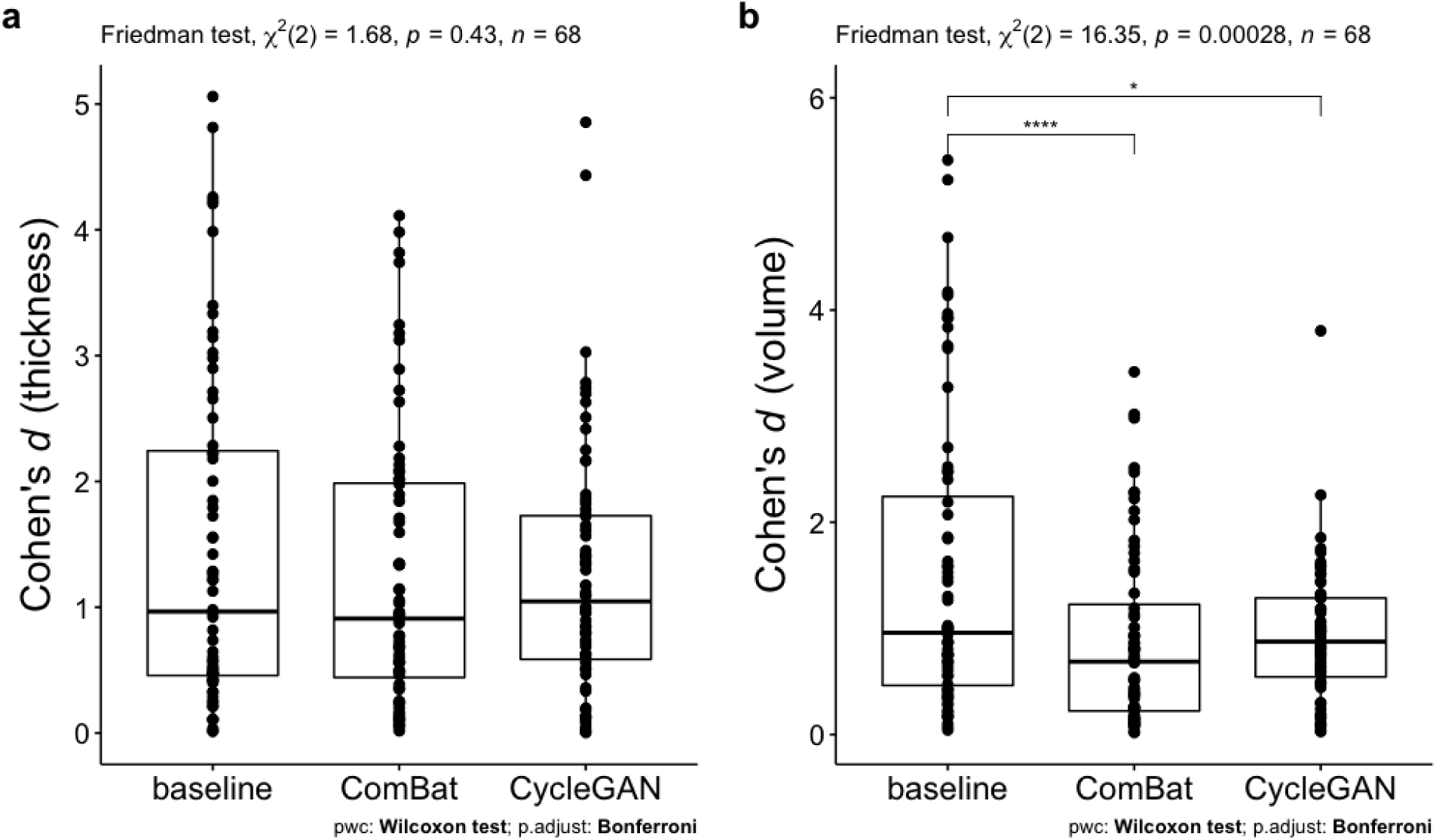
Cohen’s *d* for cortical thickness and volume. Each dot corresponds to 68 regions and the value of each region is the average of all subjects. *, p ˂ 0.05; ****, p < 0.000005

The results for volume are shown in Figure 4b. The median effect size at baseline was 0.95 (IQR, 1.78). After applying ComBat it was 0.69 (IQR, 1.00), and after applying CycleGAN it was 0.88 (IQR, 0.74). The Friedman test showed that Cohen’s *d* was significantly different between the conditions ( (2) =16.35, p = 0.00028). Pairwise Wilcoxon signed rank tests between conditions revealed statistically significant differences in PSNR between baseline and ComBat (p = 0.000002), and baseline and CycleGAN (p = 0.028). Compared with baseline, Cohen’s *d* was significantly lower with both ComBat and CycleGAN. There was no significant difference in mean values between ComBat and CycleGAN (Wilcoxon signed rank test, p = 0.62). Both the ComBat and CycleGAN methods demonstrated similar performance under the conditions of this study.

## Discussion

In this study, we employed the CycleGAN framework to address inter-site differences in T1-weighted images and evaluated its effectiveness for harmonizing TS data. The utilization of TS data offers accurate assessment of site differences because of the incorporation of images of the same subject acquired at multiple sites. Through analyses of both image-level measures (SSIM and PSNR) and feature-level measures (cortical thickness and volume), our study demonstrated the advantages of our proposed CycleGAN method in comparison with the standard ComBat approach.

SSIM and PSNR serve well as performance metrics for assessing the quality of harmonized T1-weighted images (Chai et al., 2020, Osman et al., 2022). The SSIM values before and after applying our proposed CycleGAN method showed no significant difference, indicating that the method had no significant impact on the structural similarity of the images. This result is reasonable considering that the TS data were obtained from the same subjects at different sites, and therefore the brain structures were identical (Tong et al., 2019). However, the PSNR values showed significant improvement after applying the CycleGAN method, suggesting that it provided effective improvement of image quality. In contrast, after applying ComBat, both the SSIM and PSNR values were lower than the baseline values. This can be attributed to the normalization requirement of ComBat, which involves transforming the data into MNI space, applying ComBat, and then returning the data to subject space for PSNR and SSIM calculations.

We also conducted a feature-level evaluation of cortical thickness and volume to further assess the effectiveness of our proposed CycleGAN method. In this evaluation, the Cohen’s *d* values for cortical thickness were not significantly different between the baseline, ComBat, and CycleGAN images. In contrast, in comparison with the baseline images, Cohen’s *d* for cortical volume was significantly lower for both ComBat and CycleGAN images. These results demonstrate that CycleGAN provided successful harmonization of inter-site differences in T1-weighted MRI images, with comparable effects to ComBat (Maikusa et al., 2021). However, analysis of the results of specific regions in the atlas revealed consistent deterioration in Cohen’s *d* values for both thickness and volume in regions such as the superior frontal, caudal middle frontal, precentral, and paracentral cortices. This observation suggests that CycleGAN training might not perform optimally on images near the vertex because of the smaller cortical surface area in these regions.

In this study, we used TS data to demonstrate the influence of harmonization on inter-site images, especially regarding the preprocessing step of normalization. Evaluation at the image level revealed a decrease of SSIM and PSNR compared with baseline in the images corrected using ComBat and subsequently transformed from MNI space to subject space. This finding is of interest and deserves attention. Previous studies consistently applied normalization before applying a harmonization method, but it is necessary to reassess the effects of normalization (Zuo et al., 2021, Tian et al., 2022). Considering this aspect, we placed an emphasis on the use of non-normalized data in the training of CycleGAN. Under this condition, we observed improvements in the metrics compared with baseline. This suggests the potential efficacy of a framework for harmonization without the need for spatial normalization, specifically in the context of the CycleGAN learning framework. By employing non-normalized data, we showed promising outcomes that imply the effectiveness of harmonization without the requirement for normalization.

In revisiting the significance of our approach, it’s crucial to emphasize the impact of utilizing a constrained dataset of 40 individuals for training our CycleGAN model to correct site effects arising from non-biological factors. This aspect of our research not only demonstrates the feasibility of achieving considerable accuracy with limited data but also represents a significant stride towards practical application in real-world settings where data scarcity is a common challenge. Furthermore, the prevention of overfitting with a modest dataset size, as evidenced through the employment of the Traveling Subjects dataset, underlines the robustness of our methodology. This dataset, comprising images of the same subjects from multiple facilities, provided a unique opportunity to validate our approach’s efficacy in a manner not previously explored in the literature. The implications of these findings extend beyond the technical achievements, suggesting a broader applicability of deep learning techniques in scenarios where large datasets are not available.

The present study has several limitations that need to be considered. First, although we reduced the preprocessing requirements compared with previous research, some preprocessing was still necessary. Specifically, we conducted voxel size normalization and standardization of image dimensions. Second, it is important to note that our validation of harmonization was limited to only two sites. In current multi-site studies, imaging data are often collected from more than two sites, rendering the proposed method insufficient for inter-site harmonization (Bento et al., 2021). Therefore, further extensions of the CycleGAN framework are necessary to achieve harmonization across multiple sites. Fortunately, CycleGAN architectures capable of accommodating multiple domains have been reported, and these could be considered for future applications.

## Conclusions

This study introduced a novel method for harmonization of inter-site difference in T1-weighted images using the CycleGAN framework and evaluation using TS data. The use of TS data proved advantageous because it allowed accurate assessment of site differences through the incorporation of images from the same subjects acquired across multiple sites. Our proposed method not only exhibited improvements in terms of whole image comparisons, but also demonstrated the ability to harmonize inter-site differences at the feature level (i.e., regional cortical thickness and volume). These findings suggest the potential of using CycleGAN for harmonization of inter-site differences requiring minimal preprocessing, with promising prospects for its application in multi-site neuroimaging studies of psychiatric and neurological disorders. Future work will focus on extending the architecture from inter-site to multi-site harmonization and further reducing the preprocessing steps by incorporating three-dimensional images as input.

## Acknowledgments

The authors gratefully acknowledge the financial support provided by the KAKENHI Grant Number JP18K07597, JP22K07574, JP21K07593, JP22K15777 and JP20H00625 from the Japan Society for the Promotion of Science (JSPS). This research was also supported by Intramural Research Grant (3-9, 4-6) for Neurological and Psychiatric Disorders of NCNP.

