## Supplemental Information S1 and S2 for "Harmonizing Inter-Site Differences in T1-Weighted Images Using CycleGAN"

**Supplementary Information**

**S1 The details of training and architecture of CycleGAN**

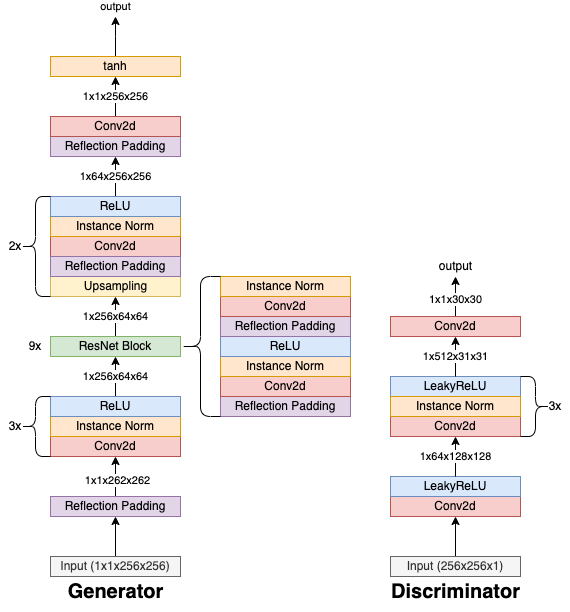

Fig. 1 Detailed architecture of CycleGAN. The size of the input corresponds to minibatch × channels × height × width, and 2× means repeat the part twice.

In optimizing our model, we established a set of hyperparameters through an extensive hyperparameter tuning process to ensure efficient and effective model training. The learning rate was set to a constant value of 0.0002 and it remains constant until the end of training. We trained the model using a mini-batch size of 1. Each 3D image was first split into axial 2D slices. All slices from all images were then used as input data.The training was conducted over 100 epochs to allow the model to learn adequately from the data without succumbing to overfitting. We employed the Adam optimizer, with beta1 and beta2 parameters set to 0.5 and 0.999, respectively, which yielded the most effective results in our preliminary trials. The cycle consistency and identity loss functions within the CycleGAN framework were weighted at 100.0 and 10.0 to balance the generative and discriminative aspects of the model.

In our architecture, the Generator is designed to generate high-quality images from a lower-resolution input. The initiation of the Generator's encoding phase entails an initial application of 3-pixel reflection padding, followed by a 2D convolutional layer with a 7-pixel kernel size and a 1-pixel stride. This convolutional layer effectively augments the feature map count from 1 to 64. Instance normalization and ReLU activation functions are applied to enhance feature representation. Subsequently, two downsampling steps are incorporated, featuring convolutional layers with a 3-pixel kernel size and 2-pixel stride, aimed at reducing spatial dimensions while enhancing feature complexity. The number of feature maps is successively increased to 128 and 256. The core of the Generator comprises multiple Residual Blocks, each consisting of two convolutional layers employing a 3-pixel kernel size and stride of 1 pixel with 256 feature maps, followed by instance normalization and ReLU activation.

Finally, the decoding phase begins with an upsampling operation to double the spatial dimensions, followed by two convolutional layers with a 3-pixel kernel size and 1-pixel stride, which decrease the feature maps to 128 and 64. The final layer employs a convolutional layer with a 7-pixel kernel size and 1-pixel stride. A hyperbolic tangent (Tanh) activation function is used to ensure that pixel values are within the range of −1 to 1.

The Discriminator, an integral component of our architectural design, plays a pivotal role in the discrimination between real and generated images. The first three convolutional layers utilize a 4-pixel kernel size, a 2-pixel stride, and a 1-pixel padding, and employ a leaky ReLU activation function. The subsequent two convolutional layers (4th and 5th layers) feature a 4-pixel kernel size, a 1-pixel stride, and a 1-pixel padding. Instance normalization and leaky ReLU activation functions are applied to enhance feature extraction and introduce non-linearity, enabling the effective differentiation between real and generated images. The Leaky ReLU parameter 'alpha' is uniformly set to 0.2 across all layers.

**S2 Model description of ComBat**

ComBat harmonization, as proposed by Fortine et al., is a popular statistical method for removing batch effects in high-dimensional data. The method assumes that feature value can be modeled as a linear model of biological variables and site effects, and that the site effects are additive or multiplicative. Therefore, the error term is modulated by an additional site-specific scaling factor. ComBat employs an empirical Bayes framework for parameter estimation, aiming to improve the estimation of the model parameters in small-sample size data.

The mathematical formulation of ComBat can be expressed as follows:

$$Y=\alpha+\beta X+\gamma+\delta\epsilon$$

Let $Y$ represent the feature value and $\alpha$ represent the average of the feature values. $\delta$ and $\gamma$ describe the additive and multiplicative site effects. $X$ is a design matrix of the covariate like age and gender, and $\beta$ is regression coefficients of design matrix.

The harmonized feature values are described as

$$Y^{ComBat}=\frac{Y-\hat{\alpha}-\hat{\beta}X-\gamma^{*}}{\delta^{*}}+\hat{\alpha}+\hat{\beta}X$$

As previously stated, the estimation of the parameters $\delta$ and $\gamma$ employs empirical Bayes.
